# Photoactivatable Blue Fluorescent Protein

**DOI:** 10.1101/2023.12.12.571285

**Authors:** Paul Gaytán, Abigail Roldán-Salgado

## Abstract

Photoactivable and photoswitchable fluorescent proteins (FPs) are sophisticated molecular tools that in combination with super-resolution microscopy are helping to elucidate many biological processes. Through the Y66H mutation in the chromophore of the violet fluorescent protein sumireF we created the first photoactivatable blue fluorescent protein (PA-BFP). This protein is rapidly activated over ordinary UV transilluminators at 302 nm or 365 nm in irreversible mode. The maximum excitation and emission wavelengths of this protein, centered at 358 nm and 445 nm, respectively, resemble the values of DAPI—the blue stain widely used in fluorescence microscopy to visualize nucleic acids in cells. Therefore, the immediate use of PA-BFP in cellular biology is clear because the technology required to follow this new genetically encoded reporter at the microscopic level has already been established. PA-BFP can potentially be used together with other photoactivatable fluorescent proteins of different colors to label multiple proteins, which can be simultaneously tracked by advanced microscopic techniques.

## Introduction

Fluorescent proteins are important molecular tools that help illuminate many biological processes and answer a plethora of questions at the microscopic level.^1^ These genetic reporters may be used to study gene expression, protein localization, protein traffic, protein fate, protein–protein interactions, sensing of metabolites, sensing of physiological conditions, and many other cellular events.^2^ The ocean contains thousands of fluorescent and chemiluminescent organisms whose fluorescent proteins or luciferases await discovery.^3^ Meanwhile, we can artificially evolve already known genes to seek protein variants with new fluorescent properties.^2d, 4^ Here, we describe the discovery and characterization of an interesting photoactivatable blue fluorescent protein (PA-BFP) derived from the recently published violet protein Sumire.^5^

## Results and Discussion

The violet fluorescent protein Sumire contains a green fluorescent protein (GFP)-type chromophore whose imidazolinone ring remains in the hydrated state, shortening the delocalization of π orbitals as compared to the ordinary GFP chromophore (Supplementary Fig. 1).^5, 6^ Consequently, Sumire displays blue-shifted excitation and emission wavelengths to 340 nm and 414 nm, respectively, taking the original GFP as a reference, whose maximum excitation and emission wavelengths are centered at 395 nm and 509 nm, respectively.^7^

Sumire was derived from superfolder GFP (sfGFP)^8^ through nine site-directed mutations: T65G, Q69A, Y145G, N146I, H148G, F165Y, T203, S205V, and V224R.^5^ However, when we assembled the gene, as described in the Methods section, following a procedure modified from the original, a fortuitous variant became violet at a faster rate than the target Sumire protein (Supplementary Fig. 2). This improved variant contained the original phenylalanine amino acid in position 165 instead of the expected tyrosine residue and was named SumireF.

Directed evolution of the SumireF gene by error-prone PCR mutagenesis^9^ and screening of the library over a UV transilluminator at 302 nm revealed the presence of a colony displaying a brilliant sky blue phenotype (Fig. 1A). DNA sequencing of the plasmid recovered from this colony confirmed that the encoded protein contained the mutation Y66H in the center of the chromophore, as inferred by a sequence comparison with the parental proteins Sumire, SumireF, and sfGFP (Fig. 1B). Notably, many engineered blue fluorescent proteins contain histidine in this position.^10^

**Figure 1.**
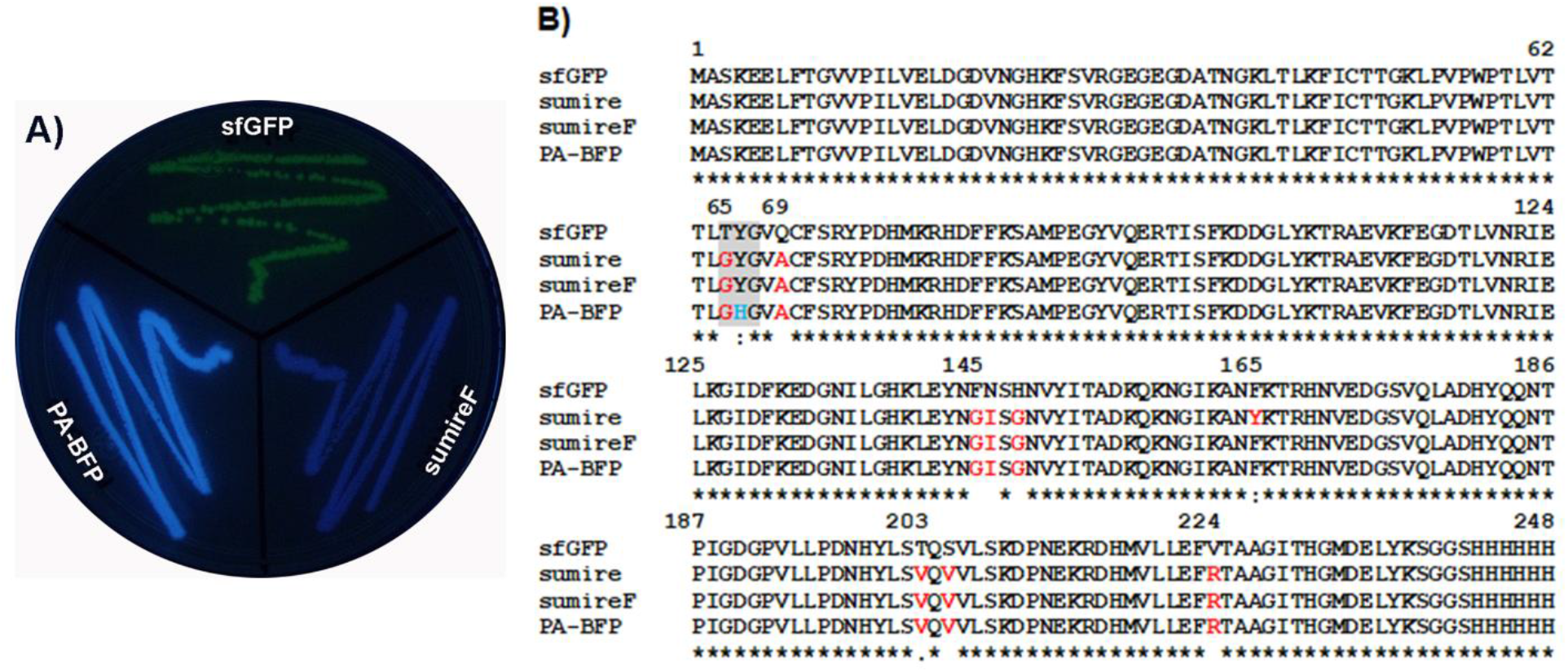
Phenotypical and sequence comparison of the photoactivatable blue fluorescent protein (PA-BFP) SumireF-Y66H. A) Comparison of *E. coli* streaks expressing the protein PA-BFP and its precursor proteins SumireF and sfGFP, visualized at 302 nm over a UV transilluminator. B) Sequence comparison of PA-BFP and its parental proteins Sumire, SumireF, and sfGFP. The chromophore-forming amino acids are shaded in gray, and the key amino acid change in PA-BFP is blue.

However, when *E. coli* cells were transformed with the plasmid harboring the mutant gene SumireF-Y66H, the resultant colonies required two minutes over the 302 nm or 365 nm UV transilluminator to fluoresce blue, as shown in the accompanying videos (PA-BFP activation at 302 nm speed 4X.mp4 and PA-BFP activation at 365 nm speed 4X.mp4). Photoactivation was also confirmed by fluorescence spectroscopy. When *E. coli* cells producing the “dark” SumireF-Y66H heterologous protein were scraped from a Petri dish, resuspended in phosphate buffer saline (PBS) buffer, and subjected to repetitive acquisition of emission spectra every two minutes, with excitation at 365 nm, the fluorescence grew in each cycle due to the cumulative fraction of activated protein, as shown in Fig. 2A. Kinetic activation of the pure dark protein, followed by fluorescence spectroscopy, demonstrated that activation at 302 nm was faster than activation at 365 nm, as shown in Fig. 2B: under 302 nm excitation, 50% of the starting “dark” protein was activated in 17.0 min, whereas 365 nm excitation produced the same ratio in 23.1 min. Complete activation of the protein, either pure or expressed in cells, is much faster on the UV transilluminator than on the fluorescence spectrometer because of the size of the excitation window (i.e., the entire screen on the transilluminator versus only 15 nm, which is the maximum excitation slit allowed in the fluorescence spectrometer at our disposal).

**Figure 2.**
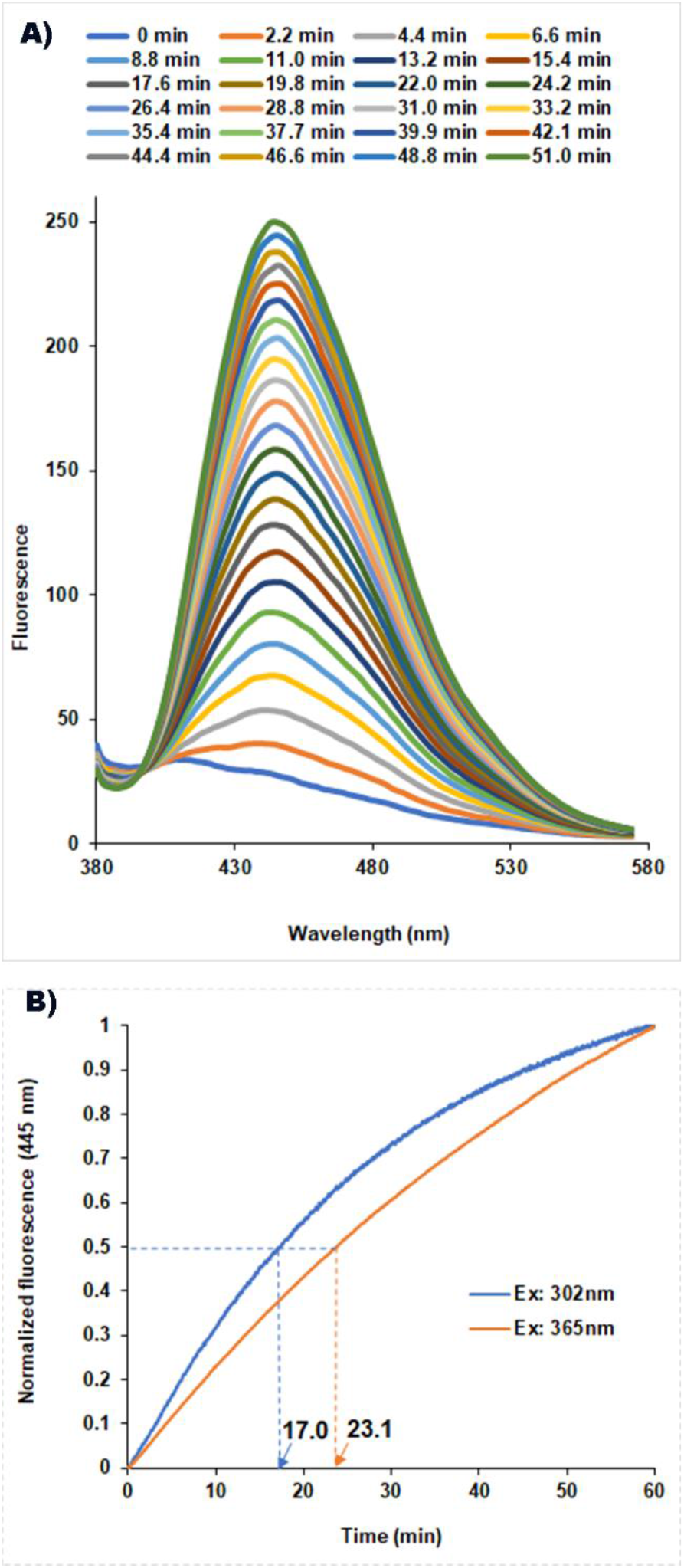
UV activation of the protein variant SumireF-Y66H (photoactivatable blue fluorescent protein [PA-BFP]). A) Repetitive acquisition of fluorescence spectra for every two minutes of *E. coli* cells resuspended in PBS buffer expressing the initially dark protein. Excitation: 360 nm; scanning rate: 100 nm/min; excitation slit: 15 nm (maximum allowed); emission slit: 5 nm. B) Photoactivation reaction of the pure dark protein PA-BFP in solution (0.13 μM) at two different excitation wavelengths, recording fluorescence at 445 nm.

From this point, the protein was named PA-BFP. There are many photoactivatable fluorescent proteins of different colors, including cyan,^11^ green,^12^ yellow,^13^ orange,^14^ and red,^15^ but PA-BFP seems to be the first photoactivatable protein in the blue range. Purification of the heterologous protein by immobilized nickel affinity chromatography yielded the “dark” protein mentioned above, whose maximum absorption was centered at 329 nm (Fig. 3A). Excitation at this wavelength produced an emission peak at 397 nm in the violet region of the visible spectrum (Fig. 3B). These blue-shifted values compared to the parental protein suggest a Sumire-like intermediate structure in which the phenol moiety of the chromophore is replaced by an imidazole ring, and the cyclized imidazolone ring remains in the hydrated form, as proposed in the post-translational mechanism shown in Fig. 4. The exposition of the intermediate protein at 365 nm over a UV transilluminator for 3 min produced a new protein version whose maximum absorbance was centered at 360 nm (Fig. 3A). Excitation at this wavelength produced an emission peak at 445 nm, as shown in Fig. 3B. Contrary to PA-BFP, the exposition of SumireF to UV did not produce any phenotypical changes to the protein.

**Figure 3.**
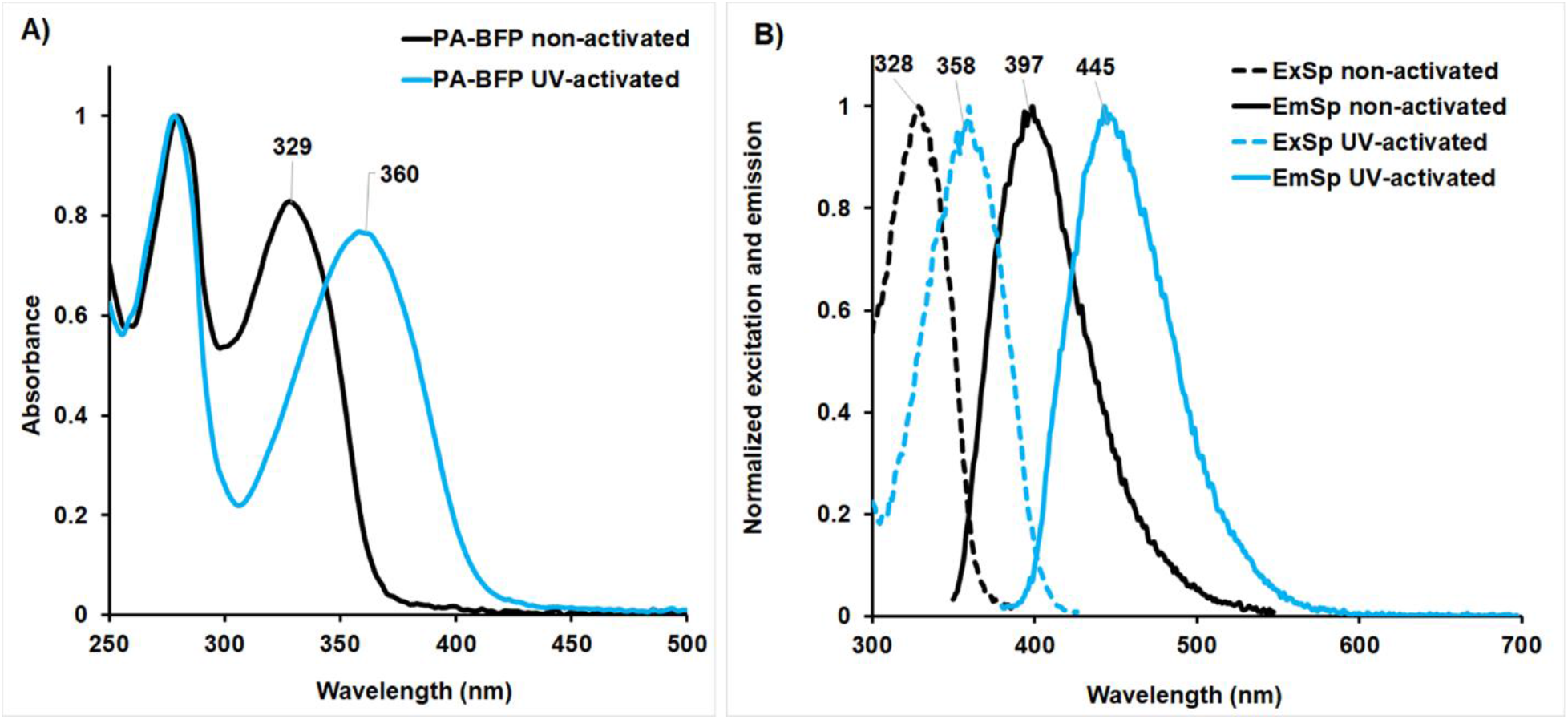
Spectroscopic analysis of photoactivatable blue fluorescent protein (PA-BFP). A) UV–Vis absorbance of the pure protein PA-BFP before and after UV activation. B) Excitation and emission spectra of the pure protein PA-BFP before and after UV activation. ExSp: excitation spectrum; EmSp: emission spectrum.

**Figure 4.**
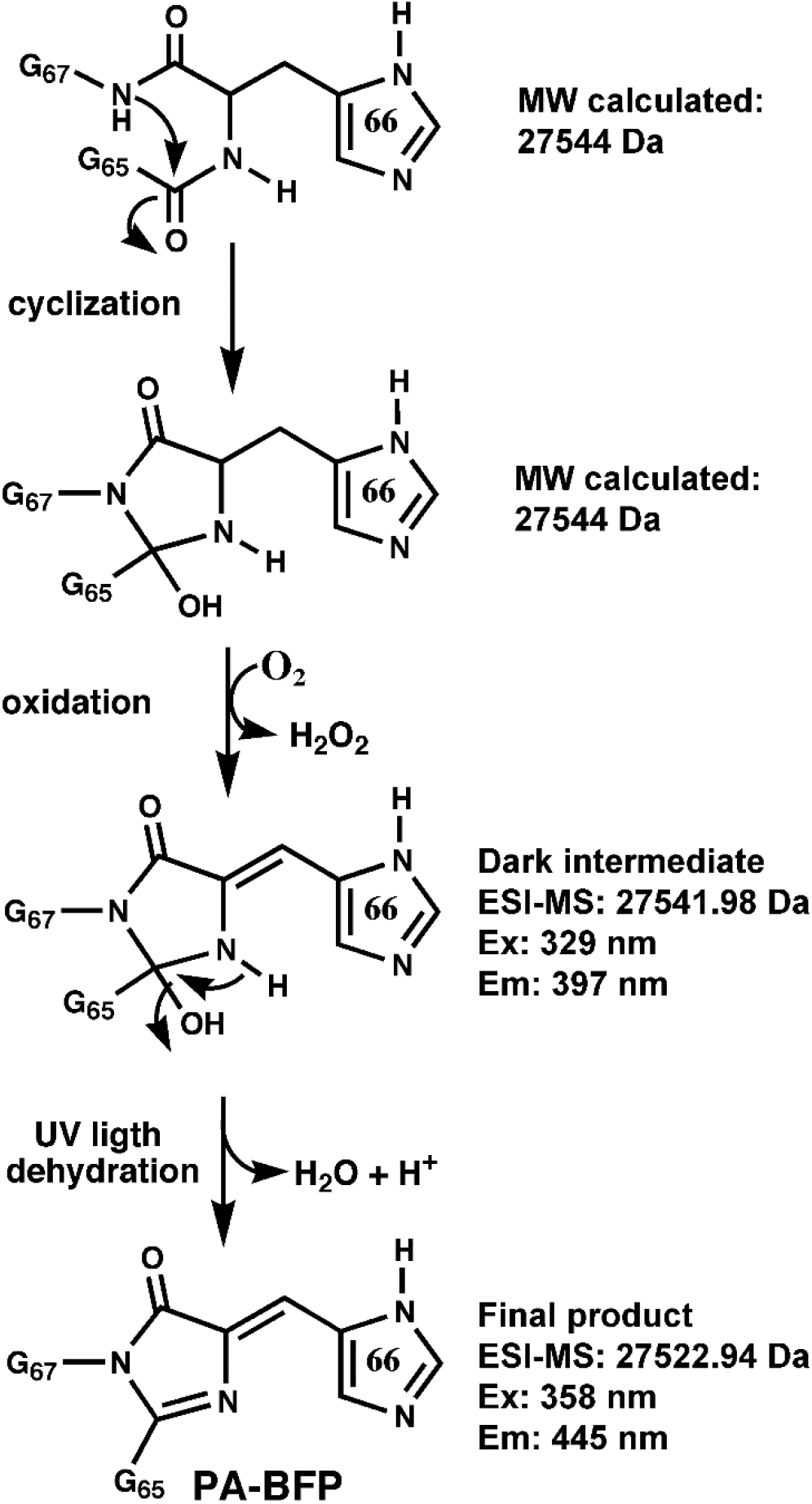
Proposed post-translational maturation mechanism of the chromophore found in the photoactivatable blue fluorescent protein (PA-BFP). The three chromophore-forming amino acids are Gly65, His66, and Gly67. Supporting results for each proposed structure are indicated on the right side. After UV activation of the dark intermediate, a molecule of water and an additional proton were eliminated.

Originally, the Sumire protein was partially designed from the photoswitchable yellow fluorescent protein (YFP) Dreiklang—a protein whose chromophore remains in the hydrated form and is activated with UV light at 365 nm to produce YFP and reverts to the dark state by irradiation with violet light at 405 nm.^16^ PA-BFP activation was irreversible, and exposing the protein to violet light did not produce a return to the dark state. Therefore, we hypothesize that PA-BFP is activated via a photolytic dehydration process as the last step in the maturation pathway of the chromophore (Fig. 4) because fluorescence emerges immediately after UV exposure. Therefore, the cyclization and oxidation steps must precede dehydration because the dark intermediate is already oxidized; it contains two fewer hydrogens than the expected unmodified polypeptide, as confirmed by electrospray ionization-mass spectrometry-liquid chromatography (ESI-MS-LC). The expected molecular weight of the unmodified polypeptide is 27544 Da, whereas the experimental value of the dark intermediate is 27541.98 Da (Supplementary Fig. 3), which has two fewer hydrogens than the complete polypeptide due to dehydrogenation of the Cα–Cβ bond of histidine 66. After UV activation, the dark intermediate lost 19 Da, whereas we expected only 18 Da, corresponding to the elimination of a water molecule. Therefore, one additional proton is lost to ionization in another part of the protein structure, or perhaps in the chromophore structure itself. Ultimately, the chromophore of PA-BFP (Fig. 4) must be the same as that present in classical BFPs derived from the original Aequorea Victoria GFP. Structural analysis of PA-BFP will likely identify the site of deprotonation and the identification of those amino acids that are important for arresting protein maturation in the oxidation step.

Sodium dodecyl sulfate polyacrylamide gel electrophoresis (SDS-PAGE) analysis of PA-BFP revealed the presence of only one band around 28 kDa after boiling the sample under denaturing conditions (Supplementary Fig. 4). This ruled out breakage of the polypeptide backbone, as occurs in the activation of kaede,^17^ EosFP,^18^ KiKGR,^19^ and Dendra^20^ and further supported photolytic dehydration as the final step in the activation of the protein.

Finally, Figure 1A clearly reveals that PA-BFP is more fluorescent than its parent protein SumireF, a qualitative result that was quantitatively confirmed by determining the optical properties of both proteins. PA-BFP exhibited a molar extinction coefficient of 13983 M^-1^cm^-1^ and a quantum yield of 0.516, whereas the results for SumireF were 23065 M^-1^cm^-1^ and 0.191, respectively. The brightness of a protein is the product of both values divided by 1,000. Therefore, the brightness of PA-BFP is 7.2 versus 4.4 for SumireF. PA-BFP was found in the first round of random mutagenesis, perhaps more mutation rounds combined with selection by fluorescence-activated cell sorting will improve its maturation efficiency and, consequently, its molar extinction coefficient to yield a more fluorescent PA-BFP.

The fluorescent properties of PA-BFP resemble those exhibited by DAPI (Exmax: 358nm, Emmax: 461 nm)—the universal dye employed to stain nucleic acids in cells.

Most fluorescence microscopes are equipped with filters for DAPI; thus, the technology has already been established for use with this new protein reporter.

In summary, we have added a new tool to the palette of photoactivated fluorescent proteins that has the appropriate properties to be used in multiple-labeling experiments and analyzed by photoactivated localization microscopy^21^ and other super-resolution microscopy techniques.^22^

## Methods

### Sumire gene construction

The Sumire gene reported by Sugiura and Nagae^5^ was assembled by site-directed mutagenesis using the sfGFP^8^ gene as a template and three pairs of oligonucleotides that grouped the expected mutations: T65G/Q69A, Y145G/N146I/H148G/F165Y, and T203/S205V/V224R. These changes were incorporated into the sfGFP gene one at a time using the overlapped PCR approach.^23^ The gene was cloned as a NdeI/BamHI insert into our constitutive pJOQ plasmid, which was previously described.^24^ For purification purposes, any insert cloned with this pair of restriction sites encodes a polyhistidine tag at the carboxy end of the target protein. The ligation reaction was used to transform the electrocompetent cells of *E. coli* MC1061.

The correct assembly of the Sumire gene was confirmed by DNA sequencing of some of the transformants that displayed a slightly violet fluorescent phenotype when the Petri dishes were analyzed over a UV transilluminator at 302 nm after incubation at 37 °C for 20 h and storage at 4 °C for 24 h. In this process of sequence confirmation, a fortuitously improved Sumire mutant was discovered. It contained all the expected mutations except F165Y. This mutant, which was more fluorescent than Sumire and did not require incubation at 4 °C to complete the process of chromophore maturation, was named SumireF.

### Generation of photoactivatable blue fluorescent protein

The SumireF gene was subjected to random PCR diversification using the GeneMorph II mutagenesis kit under the conditions suggested by the manufacturer to produce 3–4 nucleotide changes per 1 Kb of the extended template.^9^ The library of mutant genes was cloned and transformed into *E. coli* cells, as described for the Sumire gene. We found a brilliant sky blue variant after screening approximately 20,000 colonies over a UV transilluminator at 302 nm.

This mutant contained the mutation Y66H localized in the center of the chromophore. *E. coli* cells were electroporated with the plasmid Sumire-Y66H and grown on solid LB/kanamycin at 37 °C for 20 h. Visualization of the Petri dishes over a UV transilluminator at 302 nm or 365 nm (benchtop 2UV transilluminator from UVP supplemented with 8W 302 nm and 365 nm UV lamps) converted the nonfluorescent colonies into intense sky blue brilliant colonies after two minutes of exposition. The new protein was named PA-BFP.

### Protein purification

To begin protein purification, 250 mL of Luria–Bertani broth containing kanamycin at 30 μg/mL was inoculated with a colony of PA-BFP, and the culture was incubated at 37 °C for 24 h under agitation at 200 rpm. Next, the cells were recovered by centrifugation and resuspended in 12 mL of PBS buffer. The suspension was cooled in ice water and sonicated twice for 2 min with intermediate cooling. The cell debris was removed by centrifugation, and the supernatant was loaded onto a purification column containing 5 g of Ni-NTA resin, which was previously equilibrated with 0.5 M NaCl in 0.1 M phosphate buffer (pH 7.0, buffer A). The stationary phase was washed with 15 mL of the same buffer, followed by 15 mL of 30 mM of imidazole in buffer A. The heterologous protein was then eluted with 15 mL of 0.3 M imidazole in buffer A. Finally, the protein was concentrated in a centrifugal filter (Amicon Ultra-15, 10K, Millipore) and washed twice with 15 ml of PBS to remove the imidazole. The protein was recovered with 1 ml of PBS buffer, and a small fraction was saved at 4 °C as a “dark” species for subsequent experiments and analyses. The remaining protein was transferred to a quartz cuvette and activated for 3 min at 365 nm over a UV transilluminator. The parental protein SumireF was purified in the same way. Both proteins—SumireF and activated PA-BFP—were analyzed by SDS-PAGE electrophoresis, revealing the presence of only one band of the expected size (28 KDa) when the sample was boiled and two bands when the sample was analyzed under semi-denaturing conditions. One of these bands corresponded to the denatured protein, whereas the other corresponded to the native, undenatured protein.

### UV–Visible spectroscopy

The UV–visible (UV–Vis) spectra of the pure protein PA-BFP in its dark and activated state dissolved in PBS, as well as SumireF, were obtained by recording their absorbance in the range 250 nm–550 nm using silica quartz cuvettes on the spectrophotometer IMPLEM.

### Fluorescence characterization

The excitation and emission spectra of the pure proteins diluted in PBS were recorded on an LS-55 fluorescence spectrometer from PerkinElmer. The emission spectrum of each protein was obtained by fixing the excitation wavelength at its maximum absorbance wavelength, determined by UV–Vis absorbance, and recording the emission in the range “excitation wavelength + 10 nm” up to 700 nm using 5-nm emission and excitation slits. The excitation spectra were recorded in a similar mode by fixing the emission wavelength at 10 nm above its maximum and recording excitation from 350 nm to the maximum emission wavelength.

### Photoactivation rate of photoactivatable blue fluorescent protein

Under magnetic stirring, 2 ml of a pure and “dark” sample of PA-BFP 0.13 μM in PBS buffer was excited at 302 nm or 365 nm, recording the fluorescence at 445 nm during 1 h on the LS-55 fluorescence spectrometer from PerkinElmer, using an excitation slit of 15 nm and an emission slit of 5 nm. This instrument comes from the factory equipped with a high-energy pulsed xenon lamp.

### Protein analysis by size exclusion chromatography

In total, 100 μL of each purified protein, at approximately 2 μg/μL, was independently analyzed on a Superdex 200 10/300 GL column from Pharmacia, using 0.1 M NaCl in 0.1 M phosphate buffer pH 7.2 as the mobile phase, at a flow rate of 0.75 ml/min. Detection was achieved by UV absorption at 280 nm.

### Molecular mass determination

The molecular weight of PA-BFP in its dark and activated states was determined using ESI-MS-LC. The sample was applied in a system comprising an EASY-nLC II nanoflow pump (Thermo Fisher Co., San Jose, CA) coupled to an LTQ-Orbitrap Velos mass spectrometer (Thermo Fisher Co., San Jose, CA) with a nano-ESI source. The molecular mass of each sample was obtained by processing the data using the automatic deconvolution algorithm (Xtract RAW File).

### Determination of the quantum yield and molar extinction coefficient

The concentration of both activated proteins—PA-BFP and SumireF—was determined by the bicinchoninic acid assay using the QuantiPro™ Kit (Sigma–Aldrich), following the supplier’s instructions, and bovine serum albumin as the standard. To determine the extinction coefficient, five dilutions of each protein were prepared in duplicate, and their absorbance was determined at their maximum absorbance wavelengths (Sumire: 342 nm, PA-BFP: 360 nm). Plotting concentration (μM) versus absorbance produces a straight line, the slope of which is the extinction coefficient in units μM^-1^cm^-1^. Each of the above dilutions was diluted 20 times with PBS buffer, and fluorescence was determined at the maximum emission wavelength (Sumire: 413 nm, PA-BFP: 445 nm), exciting each protein at its maximum absorption wavelength. As a reference for determining the quantum yield, we used pure enhanced GFP (EGFP), whose maximum excitation and emission wavelengths were centered at 490 nm and 517 nm, respectively. The reported quantum yield for EGFP is 0.60.^25^

## Supporting information

PA-BFP supplemental figures

## Acknowledgments

Technical assistance from Leopoldo Güereca, Eugenio López-Bustos, Jorge Arturo Yáñez Ponce de León, Santiago Becerra-Ramírez, Lorena Hernández-Orihuela, Fernando Zamudio, Humberto Flores-Soto, and Edith Bernabé-Pérez is highly appreciated. We are especially grateful to Dr. Takuya Nishigaki and Dr. Gloria Saab for their critical review of our manuscript.

