## Supplementary material for "Photoactivatable Blue Fluorescent Protein": PA-BFP supplemental figures

Title:

**Supplementary Figures**

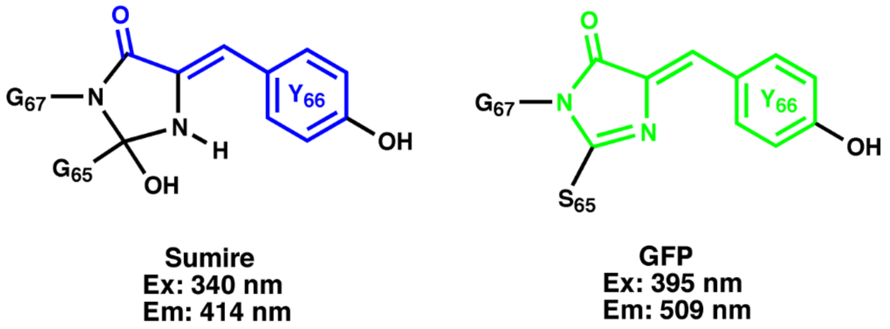


**Supplementary Figure 1.** Chromophore structures found in the proteins Sumire and avGFP. Their maximum excitation and emission wavelengths are indicated below the structures. The bonds involved in the system conjugation of π orbitals are shown in blue and green.

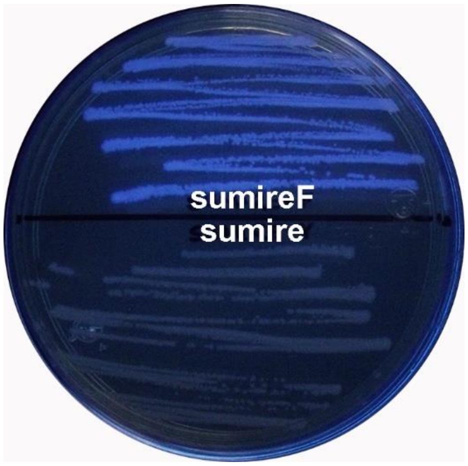


**Supplementary Figure 2**. Phenotypical comparison of the expected Sumire protein containing tyrosine at position 165, whereas the fortuitous version contains phenylalanine. This variant was named SumireF. The Petri dish was incubated at 37 °C for 20 h and then refrigerated for 24 h to enhance the maturation of the chromophore in Sumire.

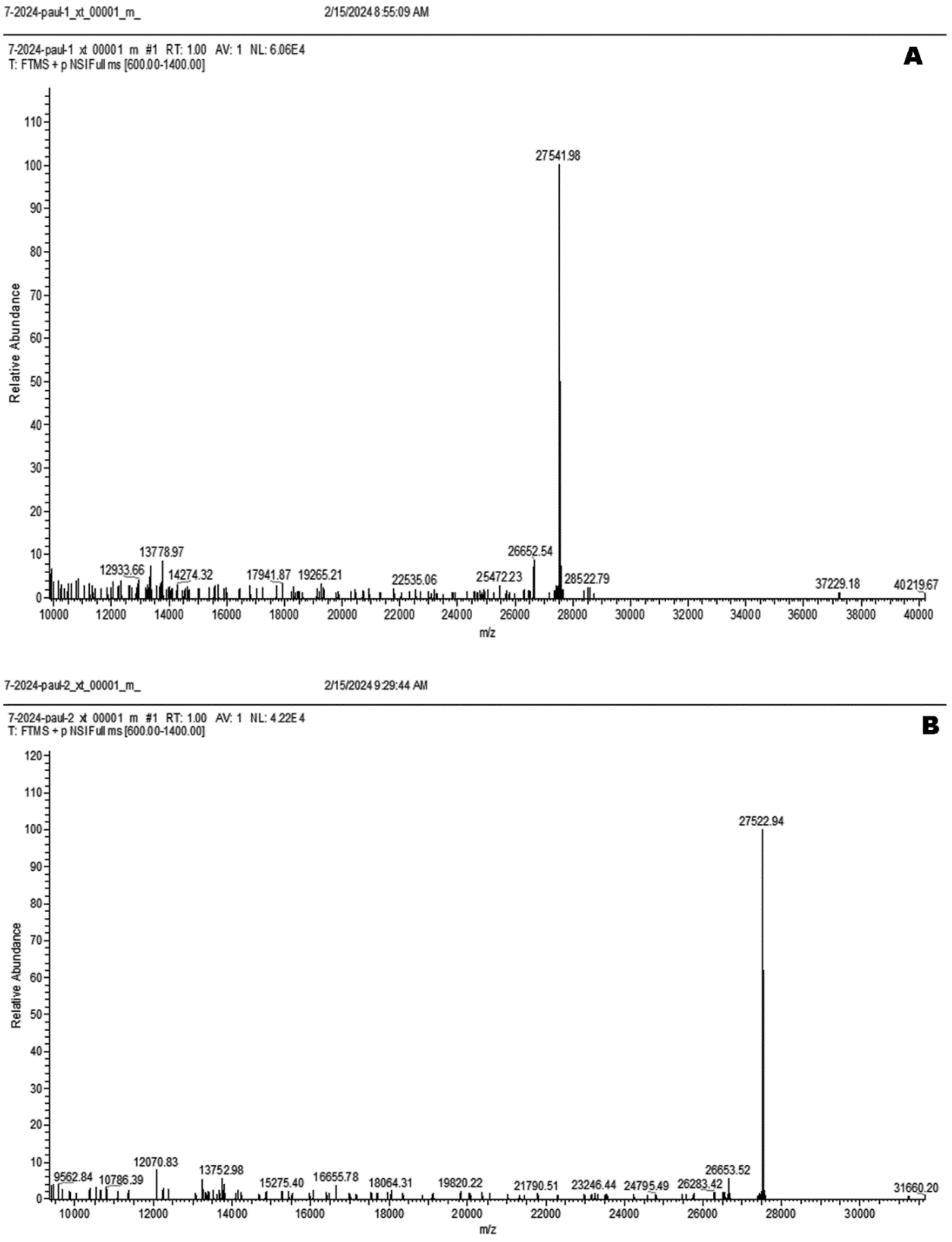


**Supplementary Figure 3.** Electrospray ionization mass spectrometry–liquid chromatography of the pure “dark” intermediate photoactivatable blue fluorescent protein (A), and the UV-activated protein (B).

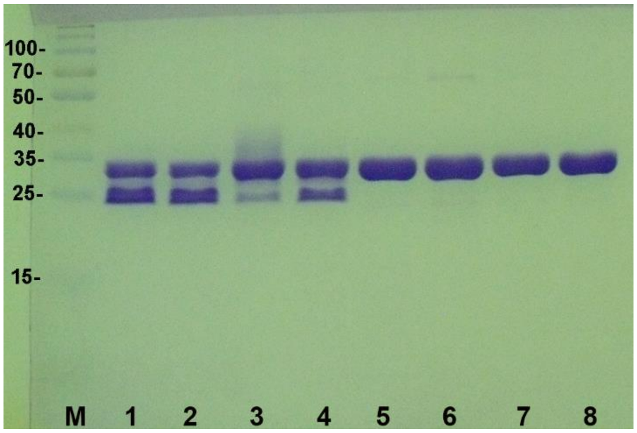


**Supplementary Figure 4**. SDS-PAGE of pure proteins. Lanes 1–4: Samples analyzed under semi-denaturing conditions (mixed with denaturing loading buffer, without boiling). Lanes 5–8: Samples analyzed under denaturing conditions (mixed with denaturing loading buffer and boiled). M: Protein marker in KDa. Lanes 1 and 5: sfGFP; lanes 2 and 6: SumireF; lanes 3 and 7: nonactivated PA-BFP; lanes 4 and 8: UV-activated PA-BFP. Analyzed protein: 5 µg per lane. Gel stained with Coomassie Blue.
